# Crimp: fast and scalable cluster relabeling based on impurity minimization

**DOI:** 10.1101/2022.03.22.485309

**Authors:** Ulrich Lautenschlager

## Abstract

**Motivation:** To analyze population structure based on multilocus genotype data, a variety of popular tools perform model-based clustering, assigning individuals to a prespecified number of ancestral populations. Since such methods often involve stochastic components, it is a common practice to perform multiple replicate analyses based on the same input data and parameter settings. Their results are typically affected by the label-switching phenomenon, which complicates their comparison and summary. Available tools allow to mitigate this problem, but leave room for improvements, in particular, regarding large input datasets.

**Results:** In this work, I present Crimp, a lightweight command-line tool, which offers a relatively fast and scalable heuristic to align clusters across multiple replicate clusterings consisting of the same number of clusters. For small problem sizes, an exact algorithm can be used as alternative. Additional features include row-specific weights, input and output files similar to those of CLUMPP (Jakobsson & Rosenberg, <u>2007</u>), and the evaluation of a given solution in terms of either CLUMPP’s and its own objective functions. Benchmark analyses show that Crimp, especially when applied to larger datasets, tends to outperform alternative tools considering runtime requirements and various quality measures.

**Availability:** Crimp’s source code along with precompiled binaries for Linux and Windows, usage guidelines and benchmark code are freely available at https://github.com/ulilautenschlager/crimp.

## Introduction

Unsupervised cluster algorithms, which aim to identify relevant groups within a set of objects (e.g., individuals or sequences), are widely used in many areas of biological data analysis. A clustering is often represented as a matrix of membership coefficients, which quantify the degree (e.g., probability or proportion) to which each clustered object (row) is assigned to each cluster (column). Typically, the identified clusters do not have meaningful labels and, in absence of ordering constraints, are listed in arbitrary order. Therefore, given a matrix of membership coefficients, each permutation of its columns represents an equivalent clustering. This ambiguity is a major reason for the incongruence of coefficient matrices across multiple clusterings of the same data, commonly referred to as ‘label-switching’ phenomenon (see Stephens, 2000; Jakobsson & Rosenberg, 2007). Since many cluster algorithms involve stochastic components, even the results of repeated runs of the same algorithm, using identical parameter settings and input data, may be affected. In order to compare or summarize a number of similar clusterings, it can be useful to align corresponding clusters across multiple clusterings. The presented program, Crimp, allows to rearrange a number of membership matrices of identical shape in order to minimize differences caused by label switching. Ideally, the remaining differences should be attributable to either noise or truly different ways of grouping the data, sometimes referred to as ‘genuine multimodality’.

An important application area of Crimp is the analysis of population structure via model-based clustering as performed by STRUCTURE (Pritchard et al., 2000) and several related methods such as Frappe (Tang et al., 2005), ADMIXTURE (Alexander et al., 2009), fastStructure (Raj et al., 2014) and MavericK (Verity & Nichols, 2016). To account for the stochasticity inherent to most of such methods, it is a common practice to perform multiple replicate analyses. Several tools are available to compare or summarize the obtained clusterings. These, as well as the presented strategy only rely on membership coefficients rather than other cluster-associated parameters. As the most widely used tool to mitigate label switching in this context, CLUMPP (Jakobsson & Rosenberg, 2007) aims to align multiple clusterings such that all pairwise similarities between membership-coefficient matrices are maximized. Popular webservers for post-processing STRUCTURE-like clusterings allow to generate input files for CLUMPP (e.g., STRUCTURE HARVESTER; Earl & von Holdt, 2012) or include it as part of their analysis pipeline (e.g., Clumpak; Kopelman et al., 2015). The latter (‘CLUMPP across K’) extends CLUMPP’s functionality to allow a comparison of clusterings comprising different numbers of clusters and to detect distinct clustering modes (i.e., to cluster the clusterings themselves) using a Markov clustering algorithm (van Dongen, 2000). pong (Behr et al., 2016), which provides functionality similar to Clumpak, implements a somewhat different approach: For each pair of membership matrices, it first infers an optimal one-to-one mapping between their corresponding columns. In case of moderate problem sizes, this can be done using an exact optimization algorithm. The distances between pairwise aligned clusterings are then used to group clusterings into modes, subject to a pre-specified distance threshold. Since, depending on the latter, there is only limited conflict among the clusterings belonging to the same mode, their clusters are simply ordered according to one randomly chosen representative clustering. This avoids to directly optimize an alignment of all clusterings, which would imply a much larger search space. Like Clumpak, pong further allows to compare clusterings that differ in their number of clusters, assuming that decreasing the latter by one can be accomodated by merging two clusters while preserving the other ones. The R package pophelper (Francis, 2016), another toolkit from this category, uses either CLUMPP or a relabeling algorithm proposed by Stephens (2000) as implemented in the R package label.switching (Papastamoulis, 2016) to align clusterings of identical size. The latter algorithm minimizes the Kullback-Leibler divergence between individual membership matrices and an averaged membership matrix through an Expectation Maximization-like (EM) algorithm.

When applying CLUMPP to larger problem instances, only its greedy heuristics, often only the rougher LargeKGreedy algorithm, are practically feasible. Since these algorithms cannot revert suboptimal decisions when building up a candidate solution, it may be difficult for them to find high quality solutions – a problem that is aggravated by large problem sizes. The cluster matching facilities provided by pong and pophelper, which both aim at a wider scope than CLUMPP, seem to be more scalable, but also lack some of CLUMPP’s functionality such as certain output options and the consideration of row-specific weights. The latter may be desirable, for instance, when clustering differently sized populations rather than individuals. The presented program, which superficially resembles CLUMPP’s functionality, aims to fill this gap and, within this limited scope, provides even better performance and scalability to large problem sizes than the aforementioned tools. The objective functions used by Crimp solely depend on impurity measures applied to rows of a single, averaged matrix of membership coefficients, which avoids a quadratic number of matrix-matrix comparisons. Crimp, which is a low-level implementation written in C, can be used as standalone command-line tool. Like CLUMPP, it can be applied in combination with existing tools for summarizing or visualizing population structure analyses or also in different contexts.

### Algorithms and implementation

The membership coefficients of a single clustering, comprising *C* objects and *K* clusters, can be written as *C* × *K* matrix, often referred to as Q-matrix. Since we deal with *R* replicate clusterings, let *c*_*ijk*_ (*i* = 1, …, *C*; *j* = 1, …, *K*; *k* = 1, …, *R*) denote the membership coefficient, which quantifies the degree to which the *i*^*th*^ object belongs to the *j*^*th*^ cluster, subject to the *k*^*th*^ clustering. Based on the individual membership coefficients, an averaged matrix (*a*_*ij*_) can be computed with 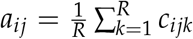. The membership coefficients are allowed to take values from [0, 1] and each row of a Q-matrix, whether individual or averaged, sums up to 1. In the following, we will effectively swap columns of the individual Q-matrices, thus these matrices and (*a*_*ij*_) will vary depending on the current state of column ordering. In fact, this is accomplished using a list of column index permutations without changing the original matrices. To simplify the notation, however, I will stick with the above notion of swapping columns. Based on the intuition that an incorrect alignment of clusters leads to a homogenization of the averaged membership coefficients, the implemented objective functions are formally based on impurity measures, namely Shannon entropy and Gini impurity. Similar to CLUMPP, optional weights *w*_*i*_ allow to control the influence of specific objects (for each *i* = 1, …, *C, w*_*i*_ >= 0). This can be useful, if, for example, the clustered objects are populations represented by a different number of sampled individuals. 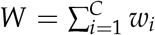 denotes the sum of these weights.

One objective function is defined as the weighted mean of the row-wise Shannon entropy of (*a*_*ij*_):

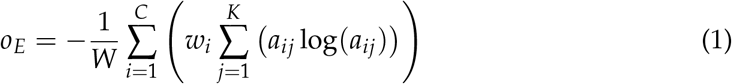

where *a*_*ij*_ log(*a*_*ij*_) := 0 if *a*_*ij*_ = 0.

Assuming equal weights *w*_*i*_ for each row index *i*, this objective function is linearly related to the total row-wise Kullback-Leibler divergence 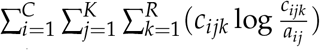tween individual Q-matrices and the averaged matrix (*a*_*ij*_) (see Supplementary Material). The latter quantity, which is often used for Stephens’ cluster relabeling method, can be expressed as *D* + *CR o*_*E*_, where 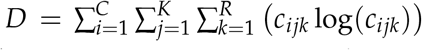 and *CR* are invariant to column swapping. Therefore, minimizing *o*_*E*_ is eqivalent to minimizing the total Kullback-Leibler divergence. However, Crimp differs from Stephens’ method by using non-EM-like algorithms. A prototypical cluster relabeling tool based on this approach has been included in AllCoPol (Lautenschlager et al., 2020) for use in a very specific context. However, the formerly used algorithm and its naive Python implementation are several orders of magnitude slower than the presented program and only suitable for relatively small problem sizes.

By default, however, Crimp minimizes the mean row-wise Gini impurity:

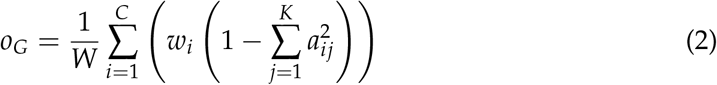

A practical advantage of using the Gini impurity over the Shannon entropy is that it does not involve logarithms and therefore allows faster computation. (2) can be interpreted in various ways (see Supplementary Material for derivations). Assuming equal weights *w*_*i*_ for simplicity, (2) can be rewritten as 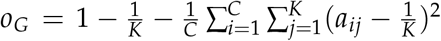,which is a linear function of the variance of the averaged membership coefficients. Therefore, minimizing *o*_*G*_ is equivalent to maximizing the variance of the averaged coefficients or, equally, minimizing the variance of matched individual coefficients, which follows from the law of total variance. For this interpretation, an alignment of all *R* Q-matrices can be viewed as a clustering of all *CKR* individual membership coefficients into *CK* clusters of size *R* each. In case of equal weights *w*_*i*_, (2) can further be expressed in terms of pairwise matrix comparison as

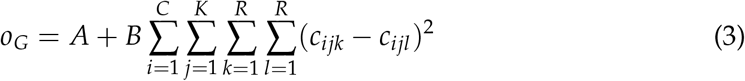

where 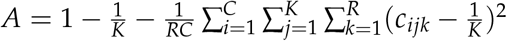 and 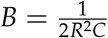 are invariant to column reordering. In other words, minimizing *o*_*G*_ minimizes the squared devitions between matched membership coefficients across all pairs of Q-matrices.

Two neighborhood-based optimization algorithms are implemented, both of which proceed by successively swapping two columns of one coefficient matrix at a time. Using auxiliary arrays to store intermediate results, this enables an evaluation of candidate solutions in *𝒪* (*C*) time because only small parts of the objective function have to be updated from solution to solution. In contrast, a naive evaluation would require *𝒪* (*CKR*) runtime per candidate solution. To avoid an accumulation of rounding errors in the course of incremental updates, the membership coefficients are internally stored using integers, providing a precision of 6 decimal places. As in CLUMPP, raw membership coefficients read from the input are normalized such that each row of a Q-matrix sums up to one.

By default, a random-restart hillclimbing heuristic is used to minimize the objective function: Starting from random column permutations, it repeatedly evaluates all possible swaps between two columns within each Q-matrix. To avoid any bias caused by the order of columns and clusterings in the input file, the order of tested swaps is randomized using a lightweight xoshiro128** (Blackman & Vigna, 2021) pseudorandom number generator. Since it would likely be inefficient to evaluate the complete swap neighborhood of given solution (i.e., 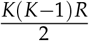 candidate solutions) before selecting the next one, each improving swap is instantly accepted. The search terminates if the current solution cannot be improved by any single swap of two columns. To mitigate its susceptibility to plateaus and local optima, this hillclimbing procedure is repeated for a predefined number of runs.

As an alternative, an exhaustive search algorithm is implemented, exploiting the fact that all permutations of a set can be generated in ways such that two consecutive permutations differ only by an interchange of two elements (for an overview of so-called bell-ringing algorithms, see Knuth, 2014). This principle can be utilized to generate all (*K*!)^*R*−1^ cluster alignments (i.e., combinations of column index permutations) such that only one swap must be evaluated for each possible solution. In contrast to the presented heuristic, this approach guarantees to find a global optimum. However, because the number of solutions to be evaluated quickly becomes prohibitive, it is only applicable to small problem sizes.

Eventually, different output files are written for the best solution found: a list of column index permutations, the matrix of averaged membership coefficients, and, optionally, the aligned individual Q-matrices. For a given solution, Crimp also allows to calculate CLUMPP’s similarity scores H and H’ although these cannot be optimized directly.

### Benchmark analyses

To compare the performance of different tools capable of aligning STRUCTURE-like Q-matrices, Crimp v1.1.0 along with CLUMPP v1.1.2, pong v1.4.7 and pophelper v2.3.0 was applied to four exemplary datasets. As biological examples, the small ‘arabid’ (*K* = 3, *C* = 95, *R* = 9) and the larger ‘chicken’ (*K* = 19, *C* = 600, *R* = 100; Rosenberg et al., 2001) datasets, both distributed with CLUMPP, were analyzed. In addition, the two simulated datasets ‘largeK’ (*K* = 100, *C* = 1000, *R* = 100) and ‘largeR’ (*K* = 25, *C* = 250, *R* = 1000) were used to demonstrate Crimp’s scalability to high values of *K* and *R*, respectively. Those are based on repeated runs of K-means clustering applied to structured random data (see kmeans_largeK.r and kmeans_largeR.r for details). As opposed to the biological datasets, the simulated datasets consist of binary Q-matrices, representing hard clusterings (partitions).

Because of differences regarding scope and output options, a fair comparison of Crimp and CLUMPP with pong and pophelper is difficult to achieve and the obtained results depend on the following decisions. Each tested program was configured to output the optimized permutations of column indices while other output was suppressed where possible. While pophelper’s alignK() function allows to rearrange Q-matrices, the applied permutations of column indices are not accessible. On the other hand, writing the aligned Q-matrices to disk using pophelpers’s clumppExport() function turned out to be very slow and may not be part of a typical pophelper workflow. To circumvent these problems, pophelper was only used to read the input matrices and the stephens() function provided by the label.switching package was then manually called with default options, as it is internally done by pophelper (see pophelper.r). Eventually, the optimized permutations were written using the R function write.table(). In the following, the described workflow will be referred to as pophelper/label.switching. In case of pong, the input matrices are partitioned into different cluster modes and separate column permutations are reported for each mode. To enforce a single mode comprising all Q-matrices, its similarity threshold parameter was set to zero. To achieve a behaviour similar to that of CLUMPP, pong was configured to use the G distance for cluster comparison. In addition, pong’s default metric, the Jaccard-Index, was used.

Each analysis was run 25 times and, for each replicate, the order of clusters and clusterings in the input was permuted. It was ensured that the measured runtimes were not distorted by memory swapping and analyses were interrupted after a maximum of 3600 seconds for the largeK dataset and 1800 seconds for the other datasets. To evaluate the obtained column index permutations, the input matrices were rearranged using an external script, thus mitigating different rounding behavior of the used tools. Crimp, which can be utilized to evaluate its input without further optimization, was then used to calculate CLUMPP’s similarity scores H and H’ as well as its own cost functions *o*_*G*_ and *o*_*E*_. Since none of the tested tools exhibited noteworthy parallelization, runtime was measured as elapsed real time. Peak memory consumption was measured as maximum resident set size. All analyses were serially performed on a Dell Optiplex 7010 desktop PC with an Intel i5-3470 CPU and 12 GB RAM. Averaged results for the chicken, largeK and largeR datasets are shown in Table 1.

**Table 1:**
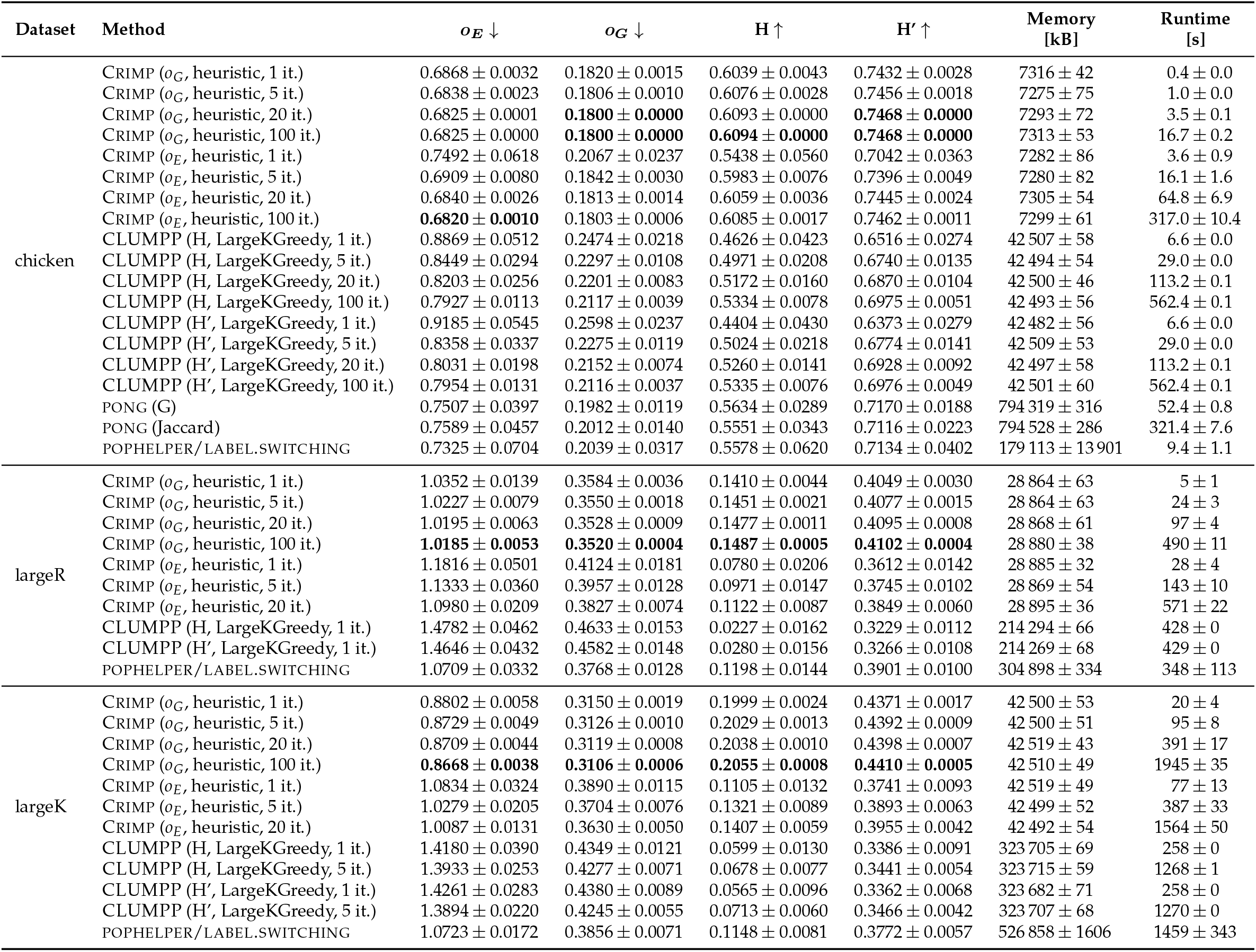
Benchmark results (mean and standard deviation) for the chicken, largeR and largeK datasets. Here, only methods for which all 25 replicate runs finished within a maximum of 1800 s (chicken, largeR) or 3600 s (largeK) are listed. As indicated by the arrows, *o*_*E*_ and *o*_*G*_ are to be minimized whereas H and H’ are to be maximized. For each dataset and qualitiy criterion, the best average results obtained are highlighted in bold.

For the small arabid dataset, only CLUMPP and Crimp were used because they are able to account for the differently sized populations. Apart from one outlier run when performing only one iteration of Crimp’s heuristic using *o*_*E*_, both programs consistently found the common global optimum of H, H’, *o*_*G*_, and *o*_*E*_, independently of the used algorithm and the optimized objective function. Runtimes ranged from less than one millisecond to about one minute (see supplementary Table S1). While, in this context, Crimp’s speed advantages are of little practical relevance, using *o*_*G*_, its exhaustive search is about two orders of magnitude faster than that of CLUMPP.

In case of the larger chicken dataset, for which either CLUMPP’s and Crimp’s exhaustive search are infeasible, each of the four evaluated scores revealed a similar pattern. Crimp using *o*_*G*_ is the fastest tool and its solutions are consistently among the best. Interestingly, optimizing *o*_*E*_ directly is not only slower due to computationally more expensive solution evaluation, but also seems to be more susceptible to local optima or plateaus as it requires a higher number of hillclimbing runs to reliably find high quality solutions. Within the allowed runtime, both pong and pophelper/label.switching are able to find solutions superior to those of CLUMPP, but less optimal than those of Crimp. As expected, pophelper/label.switching performs best in terms of the mean entropy, but even from this perspective, Crimp using *o*_*G*_ yields better scores in shorter time. In contrast, CLUMPP’s largeKgreedy algorithm leads to considerably worse scores, at least within the allowed number of at most 100 greedily constructed solutions. For 30,000 greedily constructed solutions, amounting to a runtime of approx. 47 h in our setting, Jakobsson & Rosenberg (2007) report a similarity score H=0.5546, which is similar to pong and pophelper/label.switching, but still noticeably lower than the values obtained by Crimp.

Applied to the large simulated datasets, pong did not finish within the imposed runtime limits. In case of the largeR dataset, Crimp using *o*_*G*_ was again the fastest option while yielding the best scores, followed by pophelper/label.switching and Crimp using *o*_*E*_. Similar behavior was observed for the largeK dataset, where minimizing *o*_*G*_ with Crimp takes 20 s to obtain *o*_*E*_=0.88 on average whereas pophelper/label.switch-ing, although directly minimizing the latter score, remains at 1.07 after 1100 s. For both datasets, CLUMPP’s largeKgreedy algorithm yields the least optimal scores.

Whichever of the four benchmark datasets is considered, on average, even a single iteration of Crimp’s heuristic using *o*_*G*_ (i.e., the fastest configuration) leads to results equal or better than all tested alternative tools. Compared to the latter, its results also tend to be more consistent as indicated by relatively low variances, in particular in case of *o*_*G*_. Besides, Crimp, followed by CLUMPP, shows the lowest memory demands.

## Discussion

In case of the datasets used for benchmarking, all considered quality measures (H, H’, *o*_*G*_, and *o*_*E*_) are strongly correlated. While Behr et al. (2016) recommend using the Jaccard index for pairwise comparisons instead, this comes at the cost of increased runtime requirements and an arbitrary threshold parameter, currently not exposed to the user. It should also be noted that enforcing a single mode in pong may be problematic because it is focused on aligning pairs of Q-matrices rather than reconciling multiple matrices simultaneously.

As demonstrated, especially when applied to larger datasets, Crimp tends to outper-form alternative tools in terms of runtime, solution quality and consistency. Its advantage in speed may become even more relevant if executed multiple times, for instance, in the course of Clumpak-like analyses. Crimp’s default objective function based on the Gini impurity, which emulates pairwise matrix comparison, not only allows faster computation than the mean Shannon entropy, but also seems to provide better convergence behavior of the implemented heuristic. However, the latter advantage might be problem-specific.

Especially in the context of repeated K-means clustering, it may be tempting to utilize the averaged membership matrix as a kind of consensus clustering. It should be noted that, for such use, additional postprocessing steps may be desirable, such as merging similar clusters or removing more or less empty clusters. For the analysis of population structure, K-values approaching 100 or above may appear uncommon at first glance. However, this magnitude is not unrealistic, for instance, when analyzing domestic animal breeds (e.g., Leroy et al., 2009; Funk et al., 2019; Papachristou et al., 2020).

Unlike Clumpak, pong and pophelper, Crimp is restricted to clusterings comprising the same number of clusters and does not provide additional functionality for mode detection and visualization. Nevertheless, its application helps to recognize real differences between clusterings, whether these may be regarded as noise or genuine multimodality. Moreover, Crimp can easily be integrated into common, CLUMPP-based workflows as a much faster and often more accurate alternative, applicable even to use cases which could not be handled satisfactorily before. Its heuristic can be controlled via a single, intuitive parameter, namely the number of hillclimbing runs to be performed.

As opposed to the implementation of Stephen’s cluster relabeling by the label.switch-ing package, Crimp’s performance does not depend on external solvers, but is based on efficient solution evaluation. As a consequence, it comes as a lighweight tool with-out dependencies beyond the C standard library. Future work may be dedicated to extended functionality or interfacing with other tools and programming languages.

## Acknowledgements

I would like to thank Elmar Lang for helpful discussion, Christoph Oberprieler for supporting this work, and Tankred Ott for his comments on the manuscript.

