## Supplementary Material for "Crimp: fast and scalable cluster relabeling based on impurity minimization"

### Additional results

| **Method** | $o_E \downarrow$ | $o_G \downarrow$ | **H** $\uparrow$ | **H′** $\uparrow$ | **Memory [kB]** | **Runtime [s]** |
|---|---|---|---|---|---|---|
| CRIMP ($o_G$, exhaustive) | 0.6234 | 0.2668 | 0.9696 | 0.9861 | 2384 | 0.51 |
| CRIMP ($o_E$, exhaustive) | 0.6234 | 0.2668 | 0.9696 | 0.9861 | 2380 | 6.10 |
| CRIMP ($o_G$, heuristic, 1 it.) | 0.6234 | 0.2668 | 0.9696 | 0.9861 | 2318 | 0.00 |
| CRIMP ($o_G$, heuristic, 5 it.) | 0.6234 | 0.2668 | 0.9696 | 0.9861 | 2337 | 0.00 |
| CRIMP ($o_G$, heuristic, 20 it.) | 0.6234 | 0.2668 | 0.9696 | 0.9861 | 2345 | 0.00 |
| CRIMP ($o_G$, heuristic, 100 it.) | 0.6234 | 0.2668 | 0.9696 | 0.9861 | 2331 | 0.00 |
| CRIMP ($o_E$, heuristic, 1 it.) * | 0.6498 | 0.2786 | 0.9326 | 0.9693 | 2340 | 0.00 |
| CRIMP ($o_E$, heuristic, 5 it.) | 0.6234 | 0.2668 | 0.9696 | 0.9861 | 2333 | 0.00 |
| CRIMP ($o_E$, heuristic, 20 it.) | 0.6234 | 0.2668 | 0.9696 | 0.9861 | 2337 | 0.01 |
| CRIMP ($o_E$, heuristic, 100 it.) | 0.6234 | 0.2668 | 0.9696 | 0.9861 | 2328 | 0.03 |
| CLUMPP (H, FullSearch) | 0.6234 | 0.2668 | 0.9696 | 0.9861 | 3928 | 72.14 |
| CLUMPP (H′, FullSearch) | 0.6234 | 0.2668 | 0.9696 | 0.9861 | 3931 | 72.22 |
| CLUMPP (H, Greedy, 1 it.) | 0.6234 | 0.2668 | 0.9696 | 0.9861 | 3794 | 0.00 |
| CLUMPP (H, Greedy, 5 it.) | 0.6234 | 0.2668 | 0.9696 | 0.9861 | 3802 | 0.01 |
| CLUMPP (H, Greedy, 20 it.) | 0.6234 | 0.2668 | 0.9696 | 0.9861 | 3808 | 0.02 |
| CLUMPP (H, Greedy, 100 it.) | 0.6234 | 0.2668 | 0.9696 | 0.9861 | 3810 | 0.09 |
| CLUMPP (H′, Greedy, 1 it.) | 0.6234 | 0.2668 | 0.9696 | 0.9861 | 3802 | 0.00 |
| CLUMPP (H′, Greedy, 5 it.) | 0.6234 | 0.2668 | 0.9696 | 0.9861 | 3798 | 0.01 |
| CLUMPP (H′, Greedy, 20 it.) | 0.6234 | 0.2668 | 0.9696 | 0.9861 | 3797 | 0.02 |
| CLUMPP (H′, Greedy, 100 it.) | 0.6234 | 0.2668 | 0.9696 | 0.9861 | 3814 | 0.09 |
| CLUMPP (H, LargeKGreedy, 1 it.) | 0.6234 | 0.2668 | 0.9696 | 0.9861 | 3801 | 0.00 |
| CLUMPP (H, LargeKGreedy, 5 it.) | 0.6234 | 0.2668 | 0.9696 | 0.9861 | 3801 | 0.00 |
| CLUMPP (H, LargeKGreedy, 20 it.) | 0.6234 | 0.2668 | 0.9696 | 0.9861 | 3799 | 0.01 |
| CLUMPP (H, LargeKGreedy, 100 it.) | 0.6234 | 0.2668 | 0.9696 | 0.9861 | 3809 | 0.03 |
| CLUMPP (H′, LargeKGreedy, 1 it.) | 0.6234 | 0.2668 | 0.9696 | 0.9861 | 3801 | 0.00 |
| CLUMPP (H′, LargeKGreedy, 5 it.) | 0.6234 | 0.2668 | 0.9696 | 0.9861 | 3818 | 0.00 |
| CLUMPP (H′, LargeKGreedy, 20 it.) | 0.6234 | 0.2668 | 0.9696 | 0.9861 | 3803 | 0.01 |
| CLUMPP (H′, LargeKGreedy, 100 it.) | 0.6234 | 0.2668 | 0.9696 | 0.9861 | 3820 | 0.04 |

Table S1: Averaged benchmark results for the arabid dataset. The rows of all Q-matrices were weighted by the corresponding populations sizes. As indicated by the arrows, $o_E$ and $o_G$ are to be minimized whereas H and H′ are to be maximized. For the method marked by an asterisk, one out of 25 replicate runs missed the common global optimum of $o_E$, $o_G$, H and H′.

### Derivations

#### Relationship between the total Kullback-Leibler divergence and $o_E$

$$\sum_{i=1}^{C}\sum_{j=1}^{K}\sum_{k=1}^{R}\left(c_{ijk}\log\left(\frac{c_{ijk}}{a_{ij}}\right)\right) = \underbrace{\sum_{i=1}^{C}\sum_{j=1}^{K}\sum_{k=1}^{R}\left(c_{ijk}\log\left(c_{ijk}\right)\right)}_{D} - \sum_{i=1}^{C}\sum_{j=1}^{K}\sum_{k=1}^{R}\left(c_{ijk}\log\left(a_{ij}\right)\right)$$

$$=^{a} D - R\sum_{i=1}^{C}\sum_{j=1}^{K}\left(a_{ij}\log\left(a_{ij}\right)\right)$$

$$=^{b} D + CR\, o_E$$

#### Alternative interpretations of $o_G$

If each weight $w_i$ is set to one, CRIMP'S objective function based on the Gini impurity can be written as follows:

$$o_G = \frac{1}{C}\sum_{i=1}^{C}\left(1-\sum_{j=1}^{K}a_{ij}^2\right)$$

$$= 1 - \frac{1}{C}\sum_{i=1}^{C}\sum_{j=1}^{K}a_{ij}^2$$

$$= 1 - \frac{1}{C}\sum_{i=1}^{C}\sum_{j=1}^{K}\left((a_{ij}-\frac{1}{K})^2 + \frac{2a_{ij}}{K} - \frac{1}{K^2}\right)$$

$$=^{c} 1 - \frac{1}{C}\left(\sum_{i=1}^{C}\sum_{j=1}^{K}(a_{ij}-\frac{1}{K})^2 + \frac{2C}{K} - \frac{C}{K}\right)$$

$$= 1 - \frac{1}{K} - \frac{1}{C}\underbrace{\sum_{i=1}^{C}\sum_{j=1}^{K}(a_{ij}-\frac{1}{K})^2}_{\frac{1}{R}SSB=\frac{1}{R}(SST-SSW)}$$

[a] Remember that $a_{ij} = 1/R\sum_{k=1}^{R} c_{ijk}$ .

[b] Here, equal, non-zero weights are assumed for $o_E$.

[c] Note that for each $i = 1, \ldots, C$, $\sum_{j=1}^{K} a_{ij} = 1$ .

The current order of columns of the $R$ Q-matrices implies a partitioning of all $CKR$ individual membership coefficients into $CK$ groups of size $R$ each, whose averages are given by $(a_{ij})$. The between-group sum of squares $SSB = R\sum_{i=1}^{C}\sum_{j=1}^{K}(a_{ij} - \frac{1}{K})^2$ as identified above can then be expressed as the difference of the total sum of squares $SST = \sum_{i=1}^{C}\sum_{j=1}^{K}\sum_{k=1}^{R}(c_{ijk} - \frac{1}{K})^2$ and the within-group sum of squares $SSW = \sum_{i=1}^{C}\sum_{j=1}^{K}\sum_{k=1}^{R}(c_{ijk} - a_{ij})^2$ :

$$
\begin{aligned}
o_G &= 1 - \frac{1}{K} - \frac{1}{CR}\sum_{i=1}^{C}\sum_{j=1}^{K}\sum_{k=1}^{R}\left((c_{ijk} - \frac{1}{K})^2 - (c_{ijk} - a_{ij})^2\right) \\
&=^{\mathrm{d}} 1 - \frac{1}{K} - \frac{1}{CR}\sum_{i=1}^{C}\sum_{j=1}^{K}\sum_{k=1}^{R}(c_{ijk} - \frac{1}{K})^2 + \frac{1}{2CR^2}\sum_{i=1}^{C}\sum_{j=1}^{K}\sum_{k=1}^{R}\sum_{l=1}^{R}(c_{ijk} - c_{ijl})^2 \\
&= A + B\sum_{i=1}^{C}\sum_{j=1}^{K}\sum_{k=1}^{R}\sum_{l=1}^{R}(c_{ijk} - c_{ijl})^2
\end{aligned}
$$

[d] Here, the following identity is used: given some real numbers $x_i$ $(i = 1, \ldots, n)$ with mean $\bar{x}$, $\sum_{i=1}^{n}\sum_{j=1}^{n}(x_i - x_j)^2 = 2n\sum_{i=1}^{n}(x_i - \bar{x})^2$ .
